# Density-dependent selection at low food levels leads to the evolution of population stability in *Drosophila melanogaster* even without a clear *r*-*K* trade-off

**DOI:** 10.1101/2022.06.04.494799

**Authors:** Neha Pandey, Amitabh Joshi

## Abstract

Density-dependent selection, especially together with *r*-*K* trade-offs, has been one of the most plausible suggested mechanisms for the evolution of population stability. However, experimental support for this explanation has been both meagre and mixed. One study with *Drosophila melanogaster* yielded no evidence for populations adapted to chronic larval crowding having also evolved greater population stability. Another study, on *D*. *ananassae*, suggested that populations adapted to larval crowding evolved both greater constancy and persistence stability, and the data also suggested an *r*-*K* trade-off in those populations, though the evidence for the latter was not conclusive. Moreover, theoretical work suggested that density-dependent selection could result in the evolution of greater population stability, even in the absence of an *r*-*K* trade-off. Here, we show that populations of *D*. *melanogaster*, selected for adaptation to larval crowding at very low food amounts per vial, evolve enhanced constancy and persistence stability. The enhanced population stability in the crowding-adapted populations seems to have evolved through the increased equlibrium size (*K*) and reduced sensitivity of realized population growth rates to density (*α*).There was no clear evidence for reduced intrinsic population growth rate (*r*) in the more stable crowding-adapted populations. Our study adds to the growing evidence in support of the hypothesis that population stability can evolve in response to density-dependent selection through the evolution of certain life-history traits that are associated with higher *K* and less negative *α*. We discuss our results in the light of previous work, and suggest that a model-free framework might be of great heuristic value in understanding the evolution of population stability through changes in the density-sensitivity of life-history traits, whether or not these changes result from density-dependent selection.

## INTRODUCTION

While the early understanding of population dynamics suggested that population size is regulated at a point equilibrium through the effect of density on demographic factors and through density-independent factors (reviewed in Kingsland 1995), a very interesting theoretical finding in the 1970s was that increase intrinsic growth rate in simple discrete-generation models of population dynamics destabilizes the equilibrium in population size, resulting in complex and unstable oscillatory dynamics, including chaos (May 1974). Since natural selection, all else being equal, could be expected to favour increased intrinsic growth rate, this finding implied that unstable dynamics might be fairly common. Subsequently, however, a synthesis of population dynamics data from several studies, including populations with high growth rates, suggested that population stability was more common than what would be expected from May’s finding (Hassel *et al* 1976, Thomas *et al* 1980, Mueller and Ayala 1981 a, Ellner and Turchin 1995), which generated interest in the mechanisms through which population stability could evolve. Early evolutionary explanations of population stability invoked direct selection for reduced maximal or intrinsic growth rate (*r* in the logistic or Ricker models) (Hansen 1992, Ebenman *et al* 1996), or group-selection for stable populations (Thomas *et al* 1980). A more plausible explanation was that stability can evolve as a by-product of selection on life-history traits (Mueller and Ayala 1981 b), especially if the selected traits happened to trade off with fecundity, thereby increasing population stability (Turelli and Petry 1980, Stokes *et al* 1988, Gatto 1993, Ebenman *et al* 1996, Prasad *et al* 2003).

In particular, Mueller and Ayala (1981 b) suggested that population stability could evolve via density-dependent selection through evolutionary reductions in intrinsic growth rate (*r*) under chronic crowding, through a trade-off with *K* (equilibrium population size). This hypothesis was based on the theory of density-dependent natural selection suggesting population density as a critical component of an organism’s ecology that mediates the relative fitness advantage of genotypes in a heterogeneous population (MacArthur 1962, MacArthur and Wilson 1967), first articulated in the context of selection in populations with cyclic population size dynamics by Elton (1927). Population genetic models of density-dependent selection, which were formalized in the 1970s-1980s (reviewed in Joshi *et al* 2001) proposed that the population growth rate will be high at low densities and will favor genotypes with higher intrinsic rates of growth (*r*-selection), whereas increased intra-specific competition under crowding would favor genotypes with higher competitive ability (*K*-selection). Moreover, a trade-off between *r* and *K* was expected such that no one genotype would have high fitness at both low and high density (MacArthur and Wilson 1967), as demonstrated later in *Paramecium* (Luckinbill 1979) and *Drosophila* (Mueller and Ayala 1981 b, Mueller *et al* 1991). Thus, density-dependent selection theory linked the evolution of life-history traits to population density such that genotypes that rapidly acquire resources and convert them into offspring are favored in *r*-selection, while those with better efficiency of resource utilization are favored in *K*-selection (reviewed by Reznick *et al* 2002).

Such differential evolution of life-history traits in response to density was supported by subsequent experimental investigations of traits evolving in low versus high-density environments (Luckinbill 1978, 1979, Mueller and Ayala 1981 b, Mueller 1988, Joshi and Mueller 1988, Mueller 1990, Mueller *et al* 1991), although empirical tests for population stability evolving in response to density-dependent selection have been very few (Mueller *et al* 2000, Dey *et al* 2012).

Empirical tests of density-dependent selection leading to the evolution of enhanced population stability involve experimental manipulation of population densities to implement contrasting selection pressures. Mueller and Ayala (1981 b; see also Mueller *et al* 1991) implemented such selection in *Drosophila* and developed ‘*r*-populations’ and ‘*K*-populations’ corresponding to low and high density rearing, and found that population growth rates at high density traded off with population growth rates at low density; population stability was not examined in these sets of populations. Subsequently, Mueller *et al* (1993) selected populations of *D. melanogaster* in low and high larval crowding conditions (UU and CU populations, respectively) and found that the crowding-adapted populations (CU) evolved life-history and other traits similar to the earlier-studied ‘*r*’ and ‘*K*’ populations (Mueller *et al* 1993, Joshi and Mueller 1996, Santos *et al* 1997, Borash and Ho 2001). Interestingly, however, Mueller *et al*’s (2000) investigation of population stability in the CUs and UUs when placed in a destabilizing food environment for many generations did not find evidence of enhanced constancy stability (sensu Grimm and Wissel 1997); persistence stability was not assessed as very large populations were used in the study and no extinctions were observed. The CUs did evolve traits that indicated increased competitive ability, but as in the case of the *K*-populations (Mueller 1990), these traits did not include the classic K-selected trait (MacArthur and Wilson 1967) of increased efficiency of food conversion to biomass (Joshi and Mueller 1996.

A subsequent experimental study on *D. ananassae* populations selected for adaptation to larval crowding (ACUs) presented the first evidence of both constancy and persistence stability evolving in response to density-dependent selection and it was suggested to have come about through an *r*-*K* trade-off, although the evidence for the latter was suggestive rather than conclusive (Dey *et al* 2012). Thus, these two studies (Mueller *et al* 2000, Dey *et al* 2012) yielded mixed results about the effects of density-dependent selection on population stability: stability did not evolve in the CUs (Mueller *et al* 2000) while it did in the ACUs (Dey *et al* 2012). The CU and ACU populations had experienced chronic larval crowding at very different combinations of egg number and food amount per rearing vial, with the ACUs being selected at very low food amounts (Nagarajan *et al* 2016). These differences in the ecology of experienced crowding had also led to the evolution of different sets of traits in the CUs and ACUs, with the CUs evolving higher larval feeding rates than controls (Joshi and Mueller 1996) whereas the ACUs were faster developing but did not differ from controls in larval feeding rate (Nagarajan *et al* 2016). In terms of effectiveness and tolerance (sensu Joshi *et al* 2001), the CUs had evolved traits indicative of greater effectiveness component (Joshi *et al* 2001) whereas ACUs had evolved traits likely to contribute to a higher tolerance component (Nagarajan *et al* 2016). Given these differences, Dey *et al* (2012) speculated that which specific traits evolve in response to larval crowding, and whether they primarily affect the effectiveness or tolerance components of competitive ability, might determine whether or not population stability evolves as a correlated response to density-dependent selection. However, this explanation for the difference in results between the studies of Mueller *et al* (2000) and Dey *et al* (2012) could only be suggestive, since the two studies differed not just in the food level and egg number combination at which they experienced larval crowding, but also in the species species and food medium used.

Consequently, in this study we investigated whether population stability has evolved in *D. melanogaster* populations adapted to larval crowding at the same combination of egg number and food amount, and food medium, as the ACU populations. The *D*. *melanogaster* populations we used share ancestry with the CU populations of Mueller et al (2000), rendering the comparison even more rigorous. We also investigated whether small differences in food amounts available to larvae in the population dynamics experiment interacted with selection because higher resource levels during the larval stage can increase pre-adult survivorship, fecundity, and decrease the sensitivity of growth rate to population density (Mueller and Huynh 1994, reviewed in Dey and Joshi 2018), all of which can in turn influence population dynamics (Mueller and Huynh 1994, Vaidya 2013).

## MATERIALS AND METHODS

We conducted population dynamics experiments, using *D. melanogaster* populations which were selected for adaptation to larval crowding, and their ancestral controls. We conducted these experiments at two different food amounts i.e. 1 mL and 1.5 mL per vial.

### Experimental populations

We used eight large outbred lab-maintained populations (four selected and four controls) of *D. melanogaster* which are maintained on a 21-day discrete generation cycle in all light (LL) environment at 25°C±1°C and at around 80 percent humidity. The control populations, **MB**_1-4_ (*<u>M</u>elanogaster* <u>B</u>aselines), are maintained at low larval density i.e. ∼ 70 eggs per ∼6 mL of cornmeal medium in 40 glass vials (per replicate population) of 2.2-2.4 cm inner diameter and 9.5 cm height. After eclosion (11^th^ day from egg lay when all flies emerge) the adult flies are transferred at once in respective Plexiglas cages (25 cm *×*20 cm *×* 15 cm^3^) containing a food plate with a wet cotton ball to keep up humidity. The selected populations, **MCU**_1-4_ (*<u>M</u>elanogaster* <u>c</u>rowded as larvae and <u>u</u>ncrowded as adults) have been selected for adaptation to larval crowding (competition at larval stage) and are maintained at a density of ∼600 eggs in 1.5 mL food per vial. The MCUs have been derived from their respective ancestral controls i.e. MBs with each subscript denoting ancestry. As opposed to MBs, MCUs are maintained in 12 glass vials at larval stage (to avoid adult crowding after being transferred to cages), and at the adult stage are collected in Plexiglas cages every day from 8^th^ day after egg lay till day 18 as the eclosion is spread out over many days due to larval crowding. The food plate is changed every alternate day till the 18^th^ day, and the wet cotton ball is changed at every alternate food change. On day 18 day adults from both sets of populations (i.e. MBs and MCUs) are given live acetic-acid yeast supplement till day 20, and are then allowed to lay eggs for around 18 hours. On day 21 from egg lay, eggs laid by these flies are collected in their respective densities (selected or control) to start the next generation. All populations are maintained at an adult density of about 1800 to 2000 adults. Full details of the origin and maintenance of these populations are given in Sarangi *et al* (2016). The MCU populations had undergone over 160 generations of selection prior to the population dynamics experiments.

### Population dynamics experiments

We conducted population dynamics experiments in destabilizing food environments (LH food regime: Mueller and Huynh 1994). A stabilizing environment, for example an HL food regime (<u>H</u>igh food for larvae and <u>L</u>ow food for adults i.e. absence of yeast: Mueller and Huynh 1994), induces stable dynamics in *Drosophila* populations due to a combination of relatively high larval survivorship and low adult fecundity; therefore, any differences in population stability in selected and control populations might not be detected. However, in the LH food regime (low food for larvae and <u>h</u>igh food for adults i.e. presence of yeast: Mueller and Huynh 1994) induces large fluctuations around the mean population size due to intense larval competition for food coupled with reduced sensitivity of fecundity to adult density due to the presence of yeast; therefore, evolved stability differences between different populations are much more likely to be detected in such environments (Mueller and Huynh 1994, Prasad *et al* 2003). Consequently, to look for any differences in population stability between MCUs and MBs, we set up a 31-generation long population dynamics experiment in a destabilizing (LH) food regime. We carried out this experiment at two food amounts at the larval stage i.e. one regime containing 1 mL and another containing 1.5 mL of cornmeal food for the larvae. Previously, 1 mL LH regime was shown to induce frequent extinctions (Vaidya 2013) in vial populations, and therefore, is useful to study evolution of persistence stability. We chose to also use a 1.5 mL LH regime as it parallels the food amount in the larval maintenance regime under selection for MCUs, and also to see whether food level within an LH regime interacted with selection.

From each of the four MB and four MCU populations, we derived 10 small vial populations each, after all MB and MC populations had been reared at low density (∼70 eggs/ 6 mL food) for one generation to eliminate maternal effects. We started each vial population with 8 mated females which were allowed to lay eggs in 1 mL or 1.5 mL food respectively, for 24 hours and labeled this generation as generation 0. We started transferring the eclosing flies from egg vials after day 8 from egg lay, to the matched adult collection vials containing around 4 mL of cornmeal food. Since fly eclosion is spread over several days due to competition at the larval stage, we transferred the eclosing flies daily from egg vials to their respective adult collection vials till day 18 from egg lay. We maintained vial correspondence between egg vials and adult collection vials to ensure the population identity and we did fly-transfers extremely carefully to avoid losing any flies to avoid introducing additional noise into the inherent dynamics. We moved adults to fresh adult collection vials every alternate till day 18 post egg lay. On day 18, we discarded the egg vials and gave a dab of live acetic-acid yeast on the adult collection vial wall to boost adult fecundity. On day 20 from egg collection, we transferred flies from the adult vials into new egg-laying vials with 1 mL or 1.5 mL food for next 16 hours to lay eggs for next-generation (after generation 0 the egg-laying window was decreased to 16 hours for all subsequent generations). Later, we transferred these adults into empty vials for the census counts of males and females after freezing. We also counted in the census any fly found dead during the 16-hour egg-laying phase.

The eggs laid by the flies in each vial became the next generation, i.e. density was not controlled. In parallel with the population dynamics experimental vials described above, we maintained a set of five backup vials per population whose maintenance was similar to the experimental vial populations except that backup vial populations were maintained at a low larval density to avoid the effects of crowding. Each generation, we randomly chose 5 females from each backup vial population to lay eggs for 16 hours in 6 mL of food to start the next backup generation, while the rest of the flies were discarded. Following Dey and Joshi (2006), we maintained these backup vials to reset the experimental populations (with 4 males and 4 females) in case of extinction (absence of even one male-female pair) in a vial population on day 20 post egg lay. We carried out the population dynamics experiments for 31 generations for both 1 and 1.5 mL food amounts and each generation was 21 days long. A total of 160 single-vial populations (2 selection regimes × 4 replicate populations × 2 food regimes × 10 single-vial populations) were, thus, censused over the 31 generation long population dynamics experiment. These 160 population size time series, along with the number of times each population went extinct over the 31 generations, constituted the primary data for further analyses.

### Population stability measures

*Constancy*: We compared constancy stability (*sensu* Grimm and Wissel 1997) in MBs and MCUs using 2 indices: coefficient of variation (CV) in population size and fluctuation index of population size (FI) for both 1 and 1.5 mL regimes separately. Coefficient of variation (CV) in population size measures population dispersion around the mean population size, scaled by mean population size (CV = standard deviation in population size/mean population size).We also measured constancy through FI which measures the mean one-step absolute change in population size, scaled by the mean population size (Dey and Joshi 2006), as

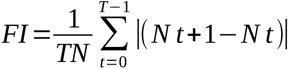

where *T* is the number of generations, *N* is the average population size, and *N_t_* and *N_t_*_+1_ are the population sizes at generations *t* and *t*+1, respectively. Constancy was interpreted as being the inverse of CV or FI, respectively.

*Persistence*: We compared persistence stability between MBs and MCUs using the frequency of extinctions in 1 and 1.5 mL regimes separately, which was calculated by dividing the number of times a population went extinct over the course of the experiment by 31 (i.e. the number of generations). Persistence was interpreted as being reflected by the inverse of the extinction probability. We counted consecutive extinctions in the same population as one extinction event because, in experiments such as these, extinctions in consecutive generations are often not independent (Dey *et al* 2008). We also calculated mean population size of each single-vial population across the 31 generations of the population dynamics experiment.

### Measuring demographic attributes

In all the single-vial populations, we examined three demographic attributes that can both respond to density-dependent selection and affect population stability: intrinsic population growth rate, equilibrium population size, and sensitivity of realized population growth rate to population density. We know from previous work that the dynamics of single-vial populations of *Drosophila* in an LH food regime are captured reasonably well by the Ricker (1954) model (Sheeba and Joshi 1998). At the same time, there is no reason to believe that the responses of *Drosophila* population dynamics to various food or selection regimes are limited by the functional form of any simple population growth model (Tung *et al* 2019, Joshi 2022). Consequently, we examined these attributes in different ways, some taking the Ricker model as the basis, while others were more empirical and model-free.

In the context of the Ricker model, we estimated the canonical parameters *r* and *K*, representing intrinsic population growth rate and equilibrium population size, respectively, as well as *α* = *r*/*K*, reflecting the sensitivity of realized population growth rate to density. These estimations were done by (a) plotting a regression line between Ln (*N_t_*_+1_/*N_t_*) on the Y-axis and *N_t_* on the X-axis, and taking the Y-intercept, X-intercept and slope as estimates of *r*, *K* and *α*, respectively, and (b) by non-linear curve fitting (following Dey *et al* 2008) using the Quasi-Newton method in Statistica vers. 5 (StatSoft 1995), followed by taking *α* = -*r*/*K*.

In addition, we also estimated realized population growth rates (*N_t_*_+1_/*N_t_*) at low (*N_t_* ≤ 30 for 1 mL food, *N_t_* < 40 for 1.5 mL food) and high (*N_t_* > 60 for 1 mL food, *N_t_*> 80 for 1.5 mL food) densities, as correlates of *r* and *K*, respectively, following the approach of Joshi *et al* (2001). We also checked the realized population growth rates at different cut-off values for low and high density to assess the robustness of the result. Finally, we estimated realized population growth rates (*N_t_*_+1_/*N_t_*) over the entire range of population densities observed during the course of the experiment, in bin sizes of 30 for 1 mL food, and 40 for 1.5 mL food.

### Statistical analyses

We used mixed model analysis of variance (ANOVA) to analyze all the response variables, with three predictor variables, two fixed, i.e. selection regime and food level, and one random, i.e. block, representing the common ancestry of each pair of MB and MCU populations with a common subscript. We performed separate ANOVAs on the coefficient of variation in population size, fluctuation index, extinction probability, average population size, intrinsic growth rate (estimated through three different methods), equilibrium population size (estimated through three different methods), and the sensitivity of realized population growth rate to population density (estimated through two methods). Separate ANOVAs on realized growth rates corresponding to different population size bins were performed for data from 1 and 1.5 mL food, because the bin sizes used differed between food levels. All analyses were performed in Statistica Ver. 5.0 (StatSoft 1995), and post-hoc comparisons used Tukey’s HSD test at *P* = 0.05.

## RESULTS

### Constancy stability

For both measures of constancy – CV and FI – the pattern of results was similar: constancy was higher in 1.5 mL than in 1 mL, and in MCUs than in MBs (Fig. 1 a, b). However, the differences between food amounts and between selection regimes were significant only in case of CV (Table 1 a), but not FI (Table 1 b). For CV, the main effects of selection and food amount were significant, but not their interaction (Table 1 a).

**Figure 1.**
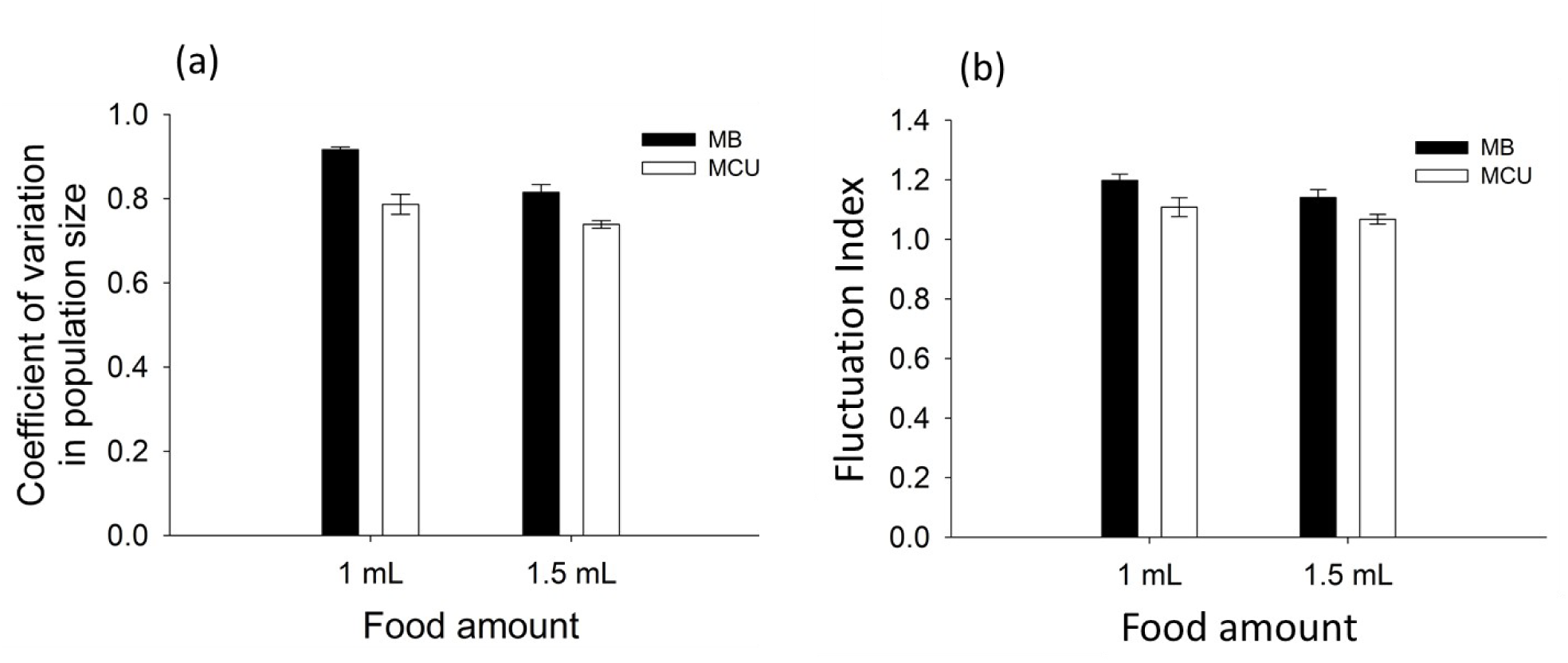
Constancy stability in MB and MCU populations in 1 and 1.5 mL food. (a) Mean coefficient of variation in population size, and (b) mean fluctuation index. Error bars around the means are standard errors based on variation among the means of the four replicate populations within each selection regime.

**Table 1:** Summary results of ANOVA done on constancy stability measured as (a) coefficient of variation in population size, and (b) fluctuation index. The table shows the main effect of selection (MCUs and MBs), food amount (1 and 1.5 mL) and their interaction. Since we were primarily interested in fixed main effects and interactions, block effects and interactions have been omitted for brevity.

| **Effect** | ***df*** | **MS** | ***df*** | **MS** | ***F*** | ***P*** |
| --- | --- | --- | --- | --- | --- | --- |
|  | **Effect** | **Effect** | **Error** | **Error** |  |  |
| a) Coefficient of variation in population size |  |  |  |  |  |  |
| Selection | 1 | 0.223 | 3 | 0.021 | 10.236 | 0.049 |
| Food amount | 1 | 0.425 | 3 | 0.007 | 60.647 | 0.004 |
| Selection × Food amount | 1 | 0.029 | 3 | 0.004 | 7.143 | 0.075 |
| b) Fluctuation index |  |  |  |  |  |  |
| Selection | 1 | 0.096 | 3 | 0.030 | 3.159 | 0.1735 |
| Food regime | 1 | 0.264 | 3 | 0.038 | 6.957 | 0.0778 |
| Selection×Food amount | 1 | 0.0028 | 3 | 0.019 | 0.141 | 0.7315 |

### Persistence stability

Persistence stability was significantly higher in MCUs than MBs, and in 1.5 mL than in 1 mL food (Fig. 2 a Table 2 a). There was also a significant interaction between selection regime and food amount (Table 2 a), driven by a much larger enhancement of persistence (much reduced extinction rate) in the MBs between 1 mL and 1.5 mL, compared to the MCUs (Fig. 2 a).

**Figure 2.**
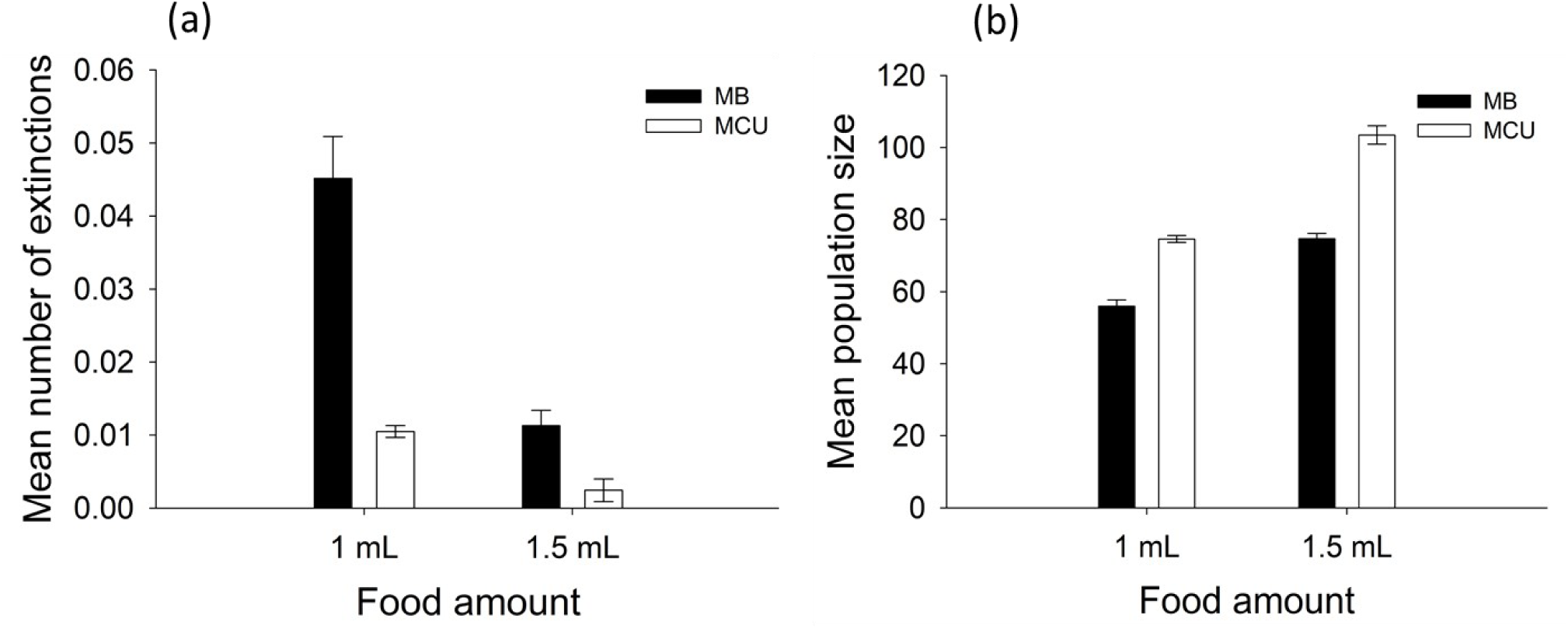
Persistence stability and average population size for MB and MCU populations in 1 mL and 1.5 mL food. (a) Mean number of extinctions per generation, and (b) mean population size. Error bars around the means are standard errors based on variation among the means of the four replicate populations within each selection regime.

**Table 2:** Summary results of ANOVA done on (a) persistence stability measured as probability of extinction, and (b) average population size. Since we were primarily interested in fixed main effects and interactions, block effects and interactions have been omitted for brevity.

| **Effect** | **df** | **MS** | **df** | **MS** | ***F*** | ***P*** |
| --- | --- | --- | --- | --- | --- | --- |
|  | **Effect** | **Effect** | **Error** | **Error** |  |  |
| a) Probability of extinction per generation |  |  |  |  |  |  |
| Selection | 1 | 0.018 | 3 | 0.000 | 42.882 | 0.007 |
| Food amount | 1 | 0.017 | 3 | 0.000 | 101.400 | 0.002 |
| Selection × Food amount | 1 | 0.006 | 3 | 0.000 | 24.000 | 0.016 |
| b) Average population size over 31 generations |  |  |  |  |  |  |
| Selection | 1 | 22492.07 | 3 | 243.008 | 92.56 | 0.002 |
| Food amount | 1 | 22653 | 3 | 63.597 | 356.19 | <0.001 |
| Selection × Food amount | 1 | 1030.2 | 3 | 104.23 | 9.88 | 0.051 |

### Mean population size

The mean population size was significantly higher in the MCUs than in the MBs, and at 1.5 mL food than 1 mL food (Fig. 2 b, Table 2 b) Going from 1 mL to 1.5 mL food, MCUs showed a greater increase in mean population size than the MBs (Fig. 2 b), but the difference was not enough to drive a significant interaction between selection regime and food amount (Table 2 b).

### Ricker-based demographic attributes

Intrinsic rate of population growth (*r*), when estimated from linear regression of Ln (*N_t+_*_1_/*N_t_*) on *N_t_*, did not differ significantly between MBs and MCUs, or between food amounts; neither the main effects of selection regime or food amount, nor their interaction, were significant (Fig. 3 a, Table 3 a). When *r* was estimated by non-linear curve fitting, there were no significant main effects of either selection regime or food amount (Table 3 b). However, the interaction between selection regime and food amount was significant (Table 3 b), and post-hoc comparisons revealed that MCUs had significantly lower estimated *r* than MBs at 1.5 mL, but not at 1 mL food amount. Apart from this one difference, not only were differences in *r* between MCUs and MBs not significant, even the magnitude of the differences was negligible (Fig. 3 a, b)

**Figure 3.**
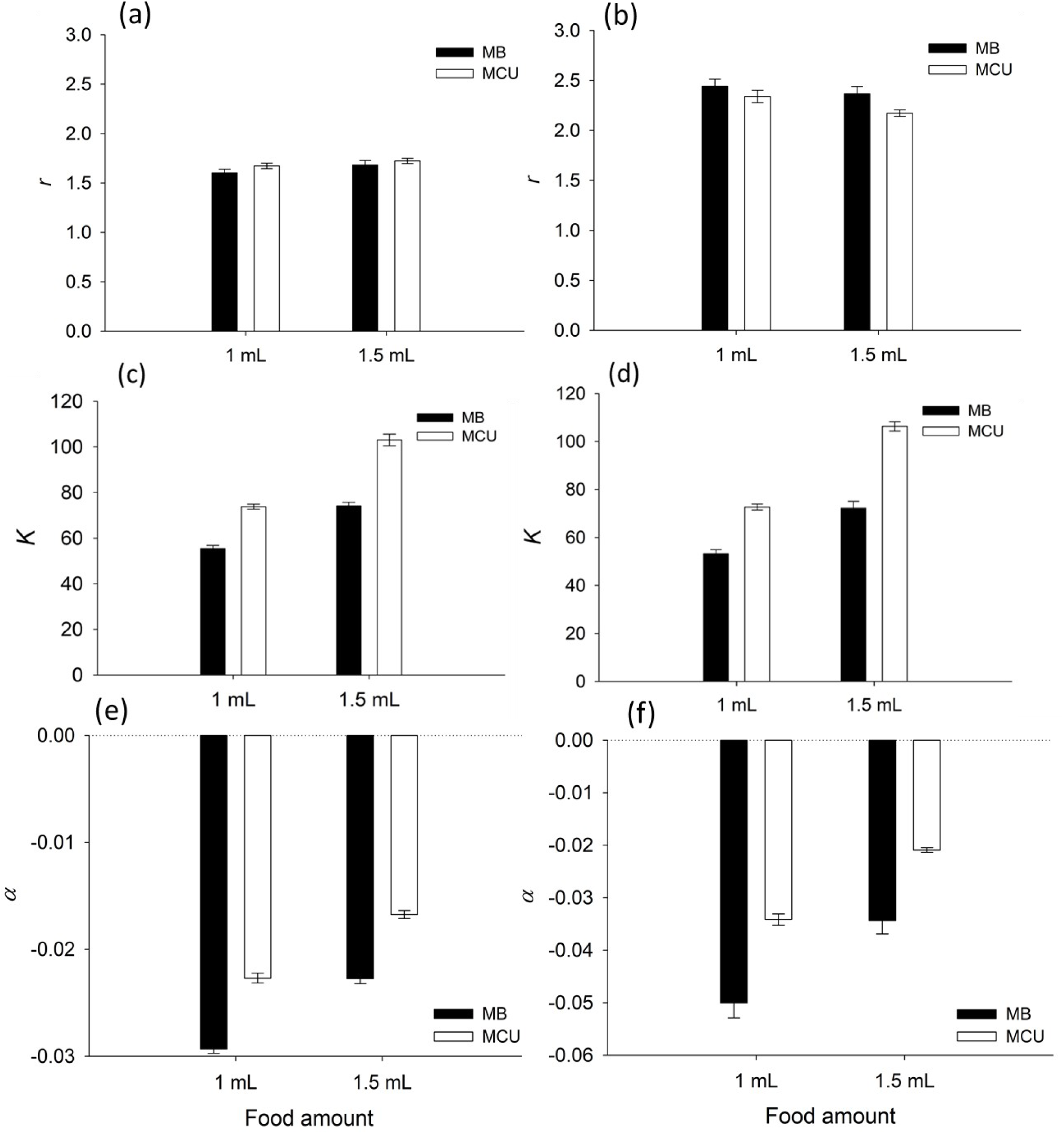
Mean demographic attributes of MB and MCU populations, based on the Ricker equation, in 1 mL and 1.5 mL food. (a) intrinsic growth rate *r* (estimated by taking the Y-intercept of the regression line between Ln (*N_t+_*_1_/*N_t_*) on Y-axis and *N_t_* on X-axis), (b) intrinsic growth rate *r* (estimated by non-linear fitting), (c) equilibrium population size *K* (estimated by taking the X-intercept of the regression line between Ln (*N_t+_*_1_/*N_t_*) on Y-axis and *N_t_* on X-axis), (d) equilibrium population size *K* (estimated by non-linear fitting), (e) sensitivity of realized growth rate to density *α* (estimated by taking the slope of the regression line between Ln (*N_t+_*_1_/*N_t_*) on Y-axis and *N_t_* on X-axis), and (f) sensitivity of realized growth rate to density *α* (estimated by non-linear fitting). Error bars around the means are standard errors based on variation among the means of the four replicate populations within each selection regime.

**Table 3:** Summary results of ANOVA done on the maximal rate of growth (*r*), based on the Ricker model, estimated from (a) linear regression of Ln (*N_t_*_+1_/*N_t_*) on *N_t_*, and (b) non-linear curve fitting. Since we were primarily interested in fixed main effects and interactions, block effects and interactions have been omitted for brevity.

| **Effect** | **df** | **MS** | **df** | **MS** | **$F$** | **$P$** |
| --- | --- | --- | --- | --- | --- | --- |
|  | **Effect** | **Effect** | **Error** | **Error** |  |  |
| a) $r$ (estimated from linear regression of $\text{Ln}(N_{t+1}/N_t)$ on $N_t$. | | | | | | |
| Selection | 1 | 0.161 | 3 | 0.062 | 2.569 | 0.207 |
| Food amount | 1 | 0.127 | 3 | 0.023 | 5.413 | 0.102 |
| Selection × Food amount | 1 | 0.008 | 3 | 0.053 | 0.149 | 0.725 |
| b) $r$ (estimated from non-linear curve fitting) | | | | | | |
| Selection | 1 | 0.527 | 3 | 0.302 | 1.740 | 0.278 |
| Food amount | 1 | 0.779 | 3 | 0.148 | 5.245 | 0.105 |
| Selection × Food amount | 1 | 0.108 | 3 | 0.005 | 18.849 | 0.022 |

The equilibrium population size (*K*) of MCUs was substantially and significantly higher than the MBs at both food amounts, and for both methods of estimation: linear regression of Ln (*N_t+_*_1_/*N_t_*) on *N_t_* and non-linear fitting (Fig. 3 c, d, Table 4 a, b). The main effect of food amount was also significant across both estimation methods (Table 4 a, b), with *K* being higher at 1.5 mL than 1 mL food for both MCUs and MBs (Fig. 3 c, d). Going from 1 mL to 1.5 mL food tended to increase *K* in the MCUs to a greater degree than in the MBs (Fig. 3 c, d), driving a significant interaction between selection regime and food amount interaction regardless of estimation method (Table 4 a, b).

**Table 4:**
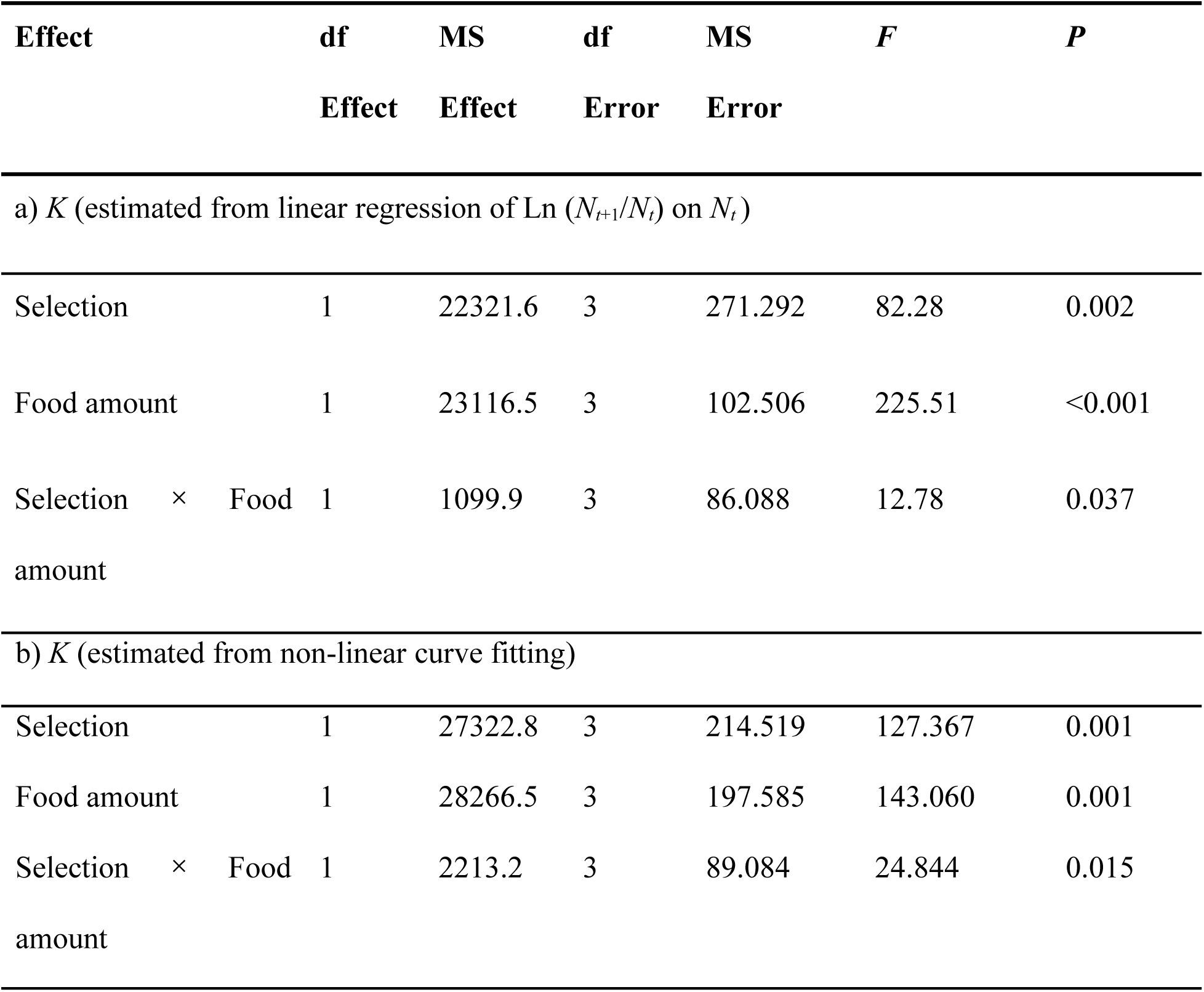
Summary results of ANOVA done on the equilibrium population size (*K*), based on the Ricker model, estimated from (a) linear regression of Ln (*N_t_*_+1_/*N_t_*) on *N_t_*, and (b) non-linear curve fitting. Since we were primarily interested in fixed main effects and interactions, block effects and interactions have been omitted for brevity.

The sensitivity of realized population growth rate to population density (*α*) showed that MCUs were significantly less sensitive to change in population density than MBs, regardless of the method of estimation (Fig. 3 e, f, Table 5 a, b). Moreover, both MCUs and MBs showed significantly reduced sensitivity (less negative values of *α*) at 1.5 mL than at 1 mL food, regardless of the method of estimation (Fig. 3 e, f, Table 5 a, b). The interaction between selection regime and food amount was not significant for either method of estimation of *α*.

**Table 5:** Summary results of ANOVA done on the sensitivity of growth rate to population density (*α*), based on the Ricker model, estimated from (a) the slope of the linear regression of Ln (*N_t_*_+1_/*N_t_*) on *N_t_*, and (b) as *α* = *r*/*K*, after non-linear curve fitting. Since we were primarily interested in fixed main effects and interactions, block effects and interactions have been omitted for brevity.

| **Effect** | **df** | **MS** | **df** | **MS** | ***F*** | ***P*** |
| --- | --- | --- | --- | --- | --- | --- |
|  | **Effect** | **Effect** | **Error** | **Error** |  |  |
| a) $\alpha$ (estimated from slope of the linear regression of $\text{Ln}(N_{t+1}/N_t)$ on $N_t$) | | | | | | |
| Selection | 1 | 0.001 | 3 | 0.000 | 285.523 | <0.001 |
| Food amount | 1 | 0.001 | 3 | 0.000 | 1752.060 | <0.001 |
| Selection × Food amount | 1 | 0.000 | 3 | 0.000 | 0.302 | 0.620 |
| b) $\alpha$ (calculated by taking a ratio $r/K$, after $r$ and $K$ were estimated by non-linear curve fitting) | | | | | | |
| Selection | 1 | 0.008 | 3 | 0.000 | 125.636 | 0.001 |
| Food amount | 1 | 0.008 | 3 | 0.000 | 31.911 | 0.010 |
| Selection × Food amount | 1 | 0.000 | 3 | 0.000 | 3.717 | 0.149 |

### Model-free demographic attributes

When we compared empirically estimated mean realized population growth rates at low versus high density in the MCUs and MBs at 1 mL and 1.5 mL food amount (Fig. 4), the only significant ANOVA effect was that of density (Table 6). Although the data suggested increased realized population growth rate in MCUs than in MBs at high density (Fig. 4), the magnitude (>25x between high and low density)) of the effect of density essentially rendered the effect of all other sources of variation on realized population growth rate relatively negligible (Table 6).

**Figure 4:**
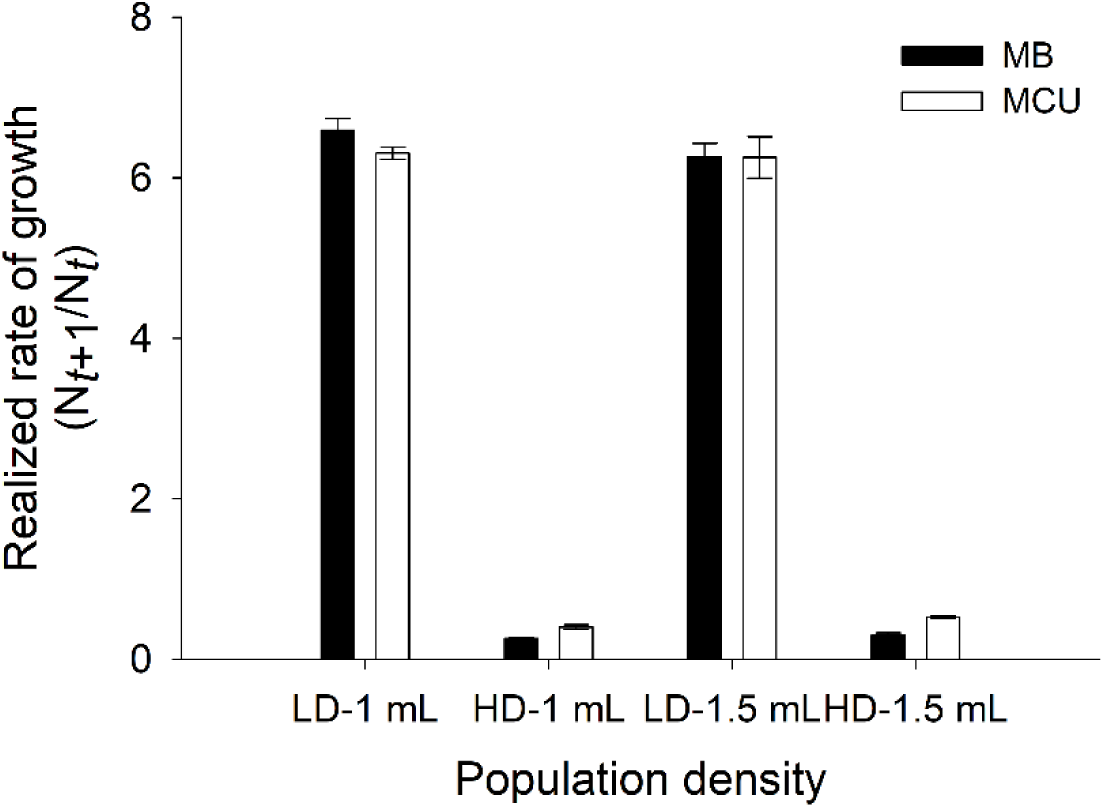
Mean empirical realized growth rates (*N_t+_*_1_/*N_t_*) at low (LD) and high density (HD)of MB and MCU populations in 1 and 1.5 mL food. LD: population size less than or equal to 30 or 40 for 1 mL and 1.5 mL, respectively. HD: population size higher than 60 or 80 for 1 mL and 1.5 mL, respectively. Error bars around the means are standard errors based on variation among the means of the four replicate populations within each selection regime.

**Table 6:**
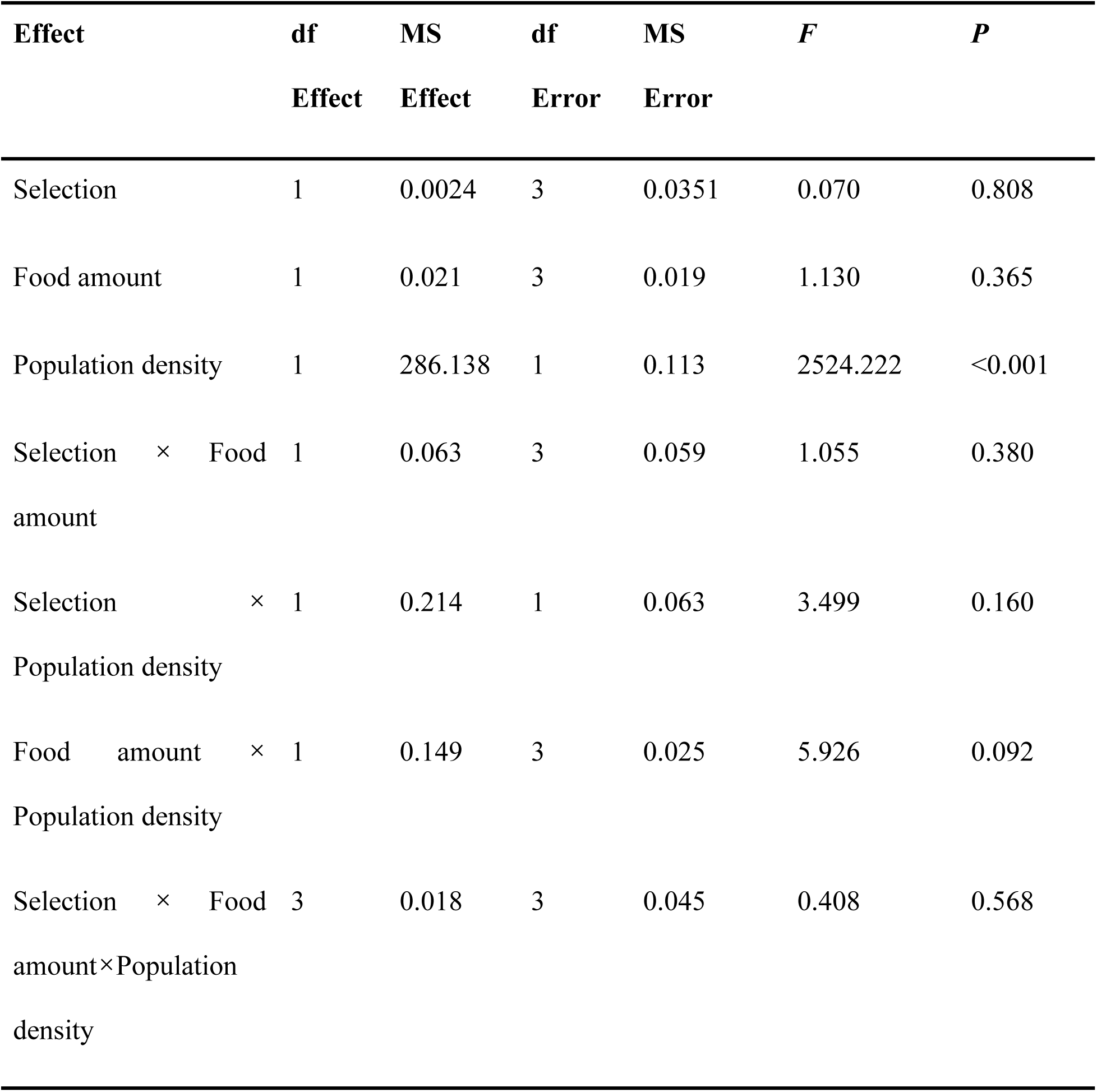
Summary results of ANOVA done on realized growth rate (*N_t_*_+1_/*N_t_*) at low and high population densities. Since we were primarily interested in fixed main effects and interactions, block effects and interactions have been omitted for brevity.

The picture became slightly clearer when we examined mean realized population growth rate across the full range of densities achieved in the single-vial populations, in bin sizes of 30 and 40 for 1 mL and 1,5 mL food, respectively (Fig. 5 a, b). There were significant ANOVA effects of selection regime, population size bin (density level), and their interaction, for mean realized population growth rate data at both 1 mL (Table 7 a) and 1.5 mL (Table 7 b) food. At both food amounts, MCUs had a higher realized population growth rate, on an average, than MBs, and realized population growth rates were substantially higher at lower population densities until a reasonably high density was attained (*N_t_* > 100 for 1 mL food, *N_t_* > 140 for 1.5 mL food), beyond which point realized population growth rates tended to level off (Fig. 5 a, b). The significant interactions, at both food amounts, between selection regime and population size bin were driven by the fact that MCUs tended to sustain significantly higher mean realized population growth rates than MBs over a range of intermediate, but not very low or very high population densities (Fig. 5 a, b). Post-hoc comparisons revealed significantly higher mean realized population growth rates in MCUs than MBs at densities between 30 and 90 individuals per vial at 1 mL food, and densities between 40 and 100 individuals per vial at 1.5 mL food (Fig. 5 a, b).

**Figure 5.**
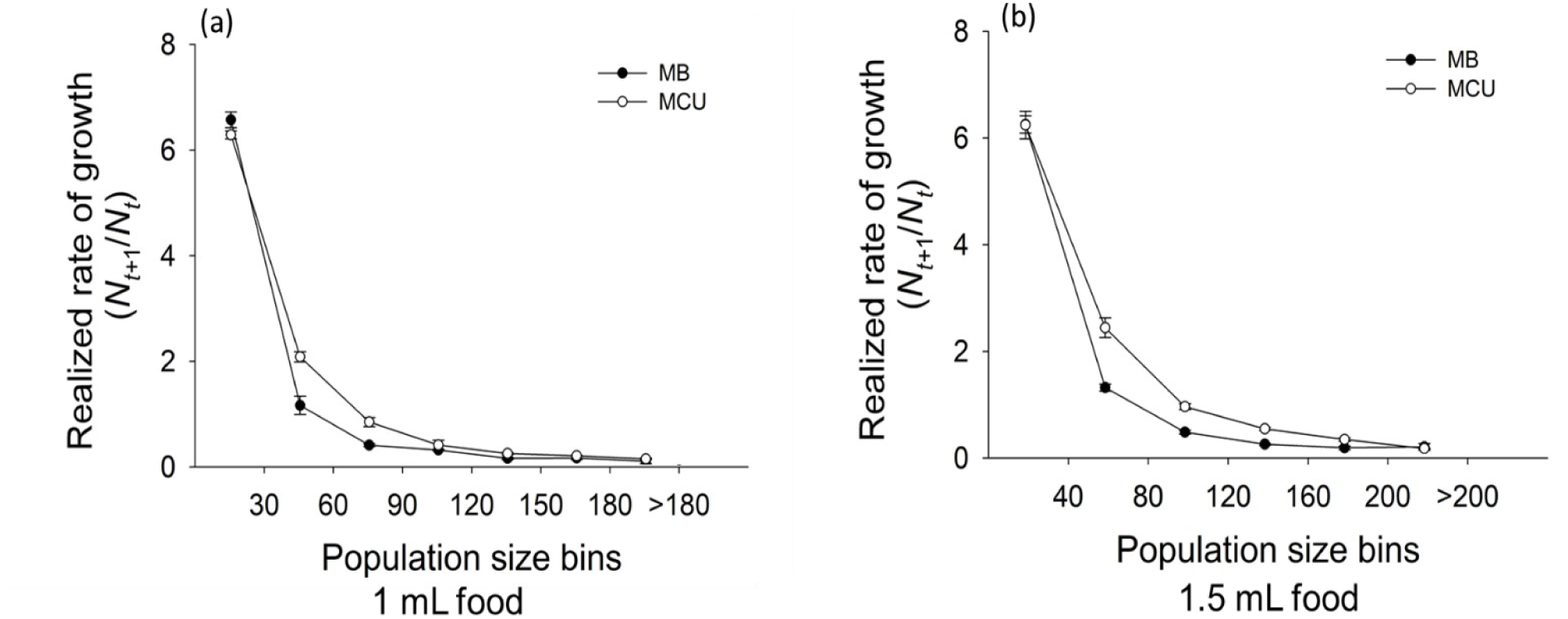
Mean empirical realized growth rates (*N_t+_*_1_/*N_t_*) at various population densities in the MB and MCU populations in (a) 1 (bin size 30 individuals) and, (b) 1.5 mL food (bin size 40 individuals). Error bars around the means are standard errors based on variation among the means of the four replicate populations within each selection regime.

**Table 7:** Summary results of ANOVA done on realized population growth rate (*N_t+_*_1_/*N_t_*) at different population density (*N_t_*) bins in MBs and MCUs in (a) 1 mL food, with a bin size of 30, and (b) 1.5 mL food, with a bin size of 40. Since we were primarily interested in fixed main effects and interactions, block effects and interactions have been omitted for brevity.

| **Effect** | **df** | **MS** | **df** | **MS** | ***F*** | ***P*** |
| --- | --- | --- | --- | --- | --- | --- |
|  | **Effect** | **Effect** | **Error** | **Error** |  |  |
| a) Realized population growth rate in 1 mL food |  |  |  |  |  |  |
| Selection | 1 | 0.515 | 3 | 0.037 | 13.858 | 0.033 |
| Bin | 6 | 41.902 | 18 | 0.035 | 1182.001 | <0.001 |
| Selection × Bin | 6 | 0.293 | 18 | 0.015 | 18.852 | <0.001 |
| b) Realized population growth rate in 1.5 mL food |  |  |  |  |  |  |
| Selection | 1 | 1.336 | 3 | 0.058 | 22.848 | 0.017 |
| Bin | 5 | 44.104 | 15 | 0.039 | 1103.469 | <0.001 |
| Selection × Bin | 5 | 0.370 | 15 | 0.047 | 7.739 | <0.001 |

## DISCUSSION

Prior to this study, two attempts to test the explanation that density-dependent selection can lead to the evolution of enhanced population stability (Mueller *et al* 2000, Prasad *et al* 2003), had yielded contradictory results. One one hand, Mueller *et al* (2000) found that populations of *D*. *melanogaster* subjected to chronic larval crowding experienced at relatively high food amounts did not evolve greater constancy than ancestral controls routinely reared at low larval density; persistence could not be compared as there were no extinctions observed in the study. On the other hand, Dey *et al* (2012) reported the evolution of greater constancy and persistence than controls in populations of *D*. *ananassae* subjected to chronic larval crowding experienced at very low food amounts. The crowding-adapted populations of Dey *et al* (2012) had evolved greater equilibrium population size, and reduced sensitivity of realized population growth rates, as compared to controls. These populations also showed considerably lower intrinsic population growth rates than controls, strongly suggestive of an *r*-*K* trade-off, but the difference was not statistically significant (Dey *et al* 2012). While it is often believed that reduced *r* is necessary for enhanced population stability, this is due to a conflation of the stability of the equilibrium population size (May 1974, Case 2000) with the stability of the observed dynamics: Dey *et al* (2012) further showed via simulations that populations could evolve greater constancy and persistence due to a higher equilibrium population size (*K*), leading to enhanced population growth rates at high densities, even in the absence of a concomitant decrease in intrinsic population growth rate at low density (*r*) due to an *r*-*K* trade-off. Dey *et al* (2012) speculated that the differences seen in these two studies with regard to the evolution, or not, of stability were likely due to the very different food amounts at which crowding-adapted populations experienced chronic larval crowding in their respective selection regimes. However, strong inferences could not be drawn because the studies of Mueller *et al* (2000) and Dey *et al* (2012) also differed in the species of *Drosophila* used. The present study was one in a set of studies designed to test the speculative hypotheses of Dey *et al* (2012) with greater rigour, using populations of *D*. *melanogaster* that shared common ancestry with those used by Mueller *et al* (2000).

Our results clearly show that *D*. *melanogaster* populations (MCU) subjected to chronic crowding at low food amounts, similar to the *D*. *ananassae* populations (ACU) of Dey *et al* (2012), also showed correlated evolution of constancy and persistence. Thus, our results strongly support the speculation of Dey *et al* (2012) that the evolution of stability in the ACUs, but not in the *D*. *melanogaster* (CU) populations of Mueller *et al* (2000), is due to differing combinations of egg number and food amount at which those two sets of populations were subjected to larval crowding. We note that the MCU populations share ancestry with the CUs, being derived from the populations that served as ancestral controls to the CUs (details in Sarangi *et al* 2016). The major difference between the MCU and CU populations is that MCUs (like the ACUs) experienced larval crowding at 600 eggs per 8 dram vial with 1.5 mL of food, whereas the CU populations had been reared at 1000-1500 eggs in 6-7 mL of food per 6-dram vial. These differences in the details of how crowding was experienced were earlier seen to result in the evolution of different sets of traits in the ACU/MCU versus the CU populations (Nagarajan *et al* 2016, Sarangi *et al* 2016, Sarangi 2018). In this context, Dey *et al* (2012) noted that the traits that evolved in the ACU populations were closer to the canonical expectation from *K*-selection, whereas traits the evolved in the CU populations were more akin to those ascribable to *α*-selection: they speculated that typical *K*-selected traits were more likely to mediate the correlated evolution of population stability, especially constancy, due to density-dependent selection than traits that evolved via *α*-selection. Our results, taken together with the findings of Sarangi *et al* (2016) and Sarangi (2018), are also consistent with the above speculation of Dey et al (2012).

In our study, while MCUs clearly had substantially greater persistence than controls (Fig. 2 a, Table 2 a), the two measures of constancy (CV and FI) gave different results: MCUs had significantly lower CV of population size than controls (Fig. 1 a, Table 1 a), but their FI values, though lower than controls on an average, did not significantly differ (Fig. 1 b, Table 1 b). The precise reason for this discrepancy is not clear at this time, but is likely to be connected to the actual distribution and sequence of population sizes in the respective time series of single-vial populations derived from the MCUs and their controls. We note that while CV reflects dispersion of population size values around the mean for a time series, the FI reflects the average one-step change in population size. Consequently, the specific sequence of population sizes in a time series can affect these two measures differently. We also note that, in Ricker-based simulations, FI does not increase monotonically with *r* beyond r ≍ 2.5 (Fig. 1 a in Sah *et al* 2013). Since the *r* values in our populations are close to that limit, at least when estimated by non-linear fitting (Fig. 3 b), it is also possible that the non-monotonic behavior of FI at high values of *r* may be playing some role here.

Compared to controls, MCUs had significantly greater mean and equilibrium population size (Figs. 2 b, 3 c, d, Tables 2 b, 4 a, b), and the values of mean and equilibrium population size were similar, suggestive of cyclic dynamics, as expected in an LH food regime (Mueller and Huynh 1994, Sheeba and Joshi 1998). There was no clear evidence of reduced intrinsic population growth rate (*r*) in the MCU populations (Figures 3 a, b, 4, 5, Table 3 a, b). Ricker based estimates of *r* showed no main effect of selection and only in 1.5 mL food was there a significantly lower *r* estimate for MCUs. Similarly, empirical estimates of realized population growth rates also indicated very similar values for MCUs and MBs at low density (Figs. 4, 5). Moreover, in the one case with the largest, and significant, difference between mean *r* in the MCUs and MBs (1.5 mL food, estimate based on non-linear fitting), the mean *r* in MBs was only about 4.6% higher than in MCUs. In the earlier study of Dey *et al* (2012), mean *r* in controls was about 12% higher than in the ACUs, even though the difference was not significant, leading the authors to conclude that there may well have been a *r*-*K* trade-off in the ACU populations, and that their study lacked the power to register it as being significant. Given the overall pattern of results for *r* in our study, we are inclined to assess the likelihood of an *r*-*K* trade-off in the MCU populations as being extremely low. We note that increased *K* could drive the observed less negative value of *α* (Fig. 3 e, f, Table 5 a, b), even in the absence of lower *r*. The large differences in Ricker-based estimates of *r* when using linear regression on log-transformed population growth rates (Fig. 3 a) versus non-linear fitting (Fig. 3 b) underscores the issues with estimating parameters of exponential functions through linearization via log-transforms pointed out by Mueller *et al* (1995).

Overall, it appears that the greater constancy and persistence of the MCUs is driven not just by higher *K* per se, but by a broader ability to maintain somewhat elevated realized population growth rates over a range of medium to high densities, even beyond *K* (Fig. 5). This observation provides empirical support for an earlier theoretical argument about how high elevated realized population growth rates around *K* can result in greater constancy as well as persistence (see Fig. 1 in Dey *et al* 2012). We note that differences like those seen between realized population growth rates of MCUs and MBs across densities (Fig. 5), while clearly indicating reduced sensitivity of growth rates to density in the MCUs, will not contribute to less negative values of *α* in the absence of differences in *r* or *K* between the two sets of populations. Thus, the pattern of differences in realized population growth rates between MCUs and MBs (Fig. 5) also suggests that the framework of simple models of population growth, like the Ricker or logistic, may not be adequate to capture how population stability changes as a result of density-dependent selection, because the pattern seen in Fig. 5 cannot be explained by changes in parameters like *r* or *K*. Generalized three-parameter versions of these models like the *θ*-logistic or *θ*-Ricker tend to capture differences in dynamics between populations with differing histories of density-dependent selection better than their canonical two-parameter counterparts (Gilpin *et al* 1976), but even these model variants cannot accommodate the possible evolution of higher realized population growth rates at densities both below and above *K*.

In terms of the effect of food level (1 mL vs 1.5 mL) in the single-vial populations in the population dynamics experiment, our findings of higher constancy and persistence stability at higher food levels are in agreement with the trend reported previously from a much shorter 10 generation study on the dynamics of MCU- and MB-derived single-vial populations on an LH food regime with 1, 2 or 3 mL of food per vial (Vaidya 2013). The stabilizing effects of higher food levels can be attributed to how the demographic attributes respond to changes in food level. Equilibrium population size (*K*), and mean population size, were higher at 1.5 mL food, whereas the sensitivity of realized population growth rate to population density (*α*) was lower. Together with Vaidya’s (2013) findings, our results suggest that increasing food levels from 1 mL to 1.5 mL per vial in a population dynamics experiment enhances stability by reducing the sensitivity of realized population growth rates to density, whereas increasing the food level further from 2 mL to 3 mL per vial does not enhance stability further. This has consequences for experimental evolution studies since the evolved differences in stability are more easily detectable at lower food levels, which should be used for studying evolutionary changes in population stability. Previously, for example, Vaidya (2013) found no difference in constancy and persistence between MCUs and MBs in a 10 generation population dynamics experiment conducted under an LH food regime with 2 mL of food per vial, even though MCUs had evolved higher *K* and less negative *α* by that time.

To conclude, our findings add to growing evidence in support of the hypothesis that population stability can evolve as a correlated response to density-dependent selection, most likely through the evolution of certain life-history traits that influence the sensitivity of fitness components to high density. Our results also support the view that both persistence and constancy can increase as a correlated response to chronic crowding even without an evolutionary reduction in intrinsic population growth rate, as long as the adaptation to crowding facilitates the maintenance of higher realized population growth rates across a range of medium to high densities. This makes it likely that density-dependent selection might be a more common contributor to the evolution of population stability than previously thought. Moreover, our results suggest that a model-free heuristic framework might be more useful than relying on simple population growth models when studying the consequences of life-history evolution for population stability, whether via density-dependent selection or not.

## ACKNOWLEDGMENTS

We thank Shruti Mallya, Sajith V. S., Satyabrata Nayak and Joy Bose for help with the population dynamics experiments, and Avani Mital, Manaswini Sarangi, N. Rajanna and Muniraju for assistance in the maintenance of selected and control populations. NP was supported by a doctoral fellowship from the Jawaharlal Nehru Centre for Advanced Scientific Research. This work was supported by a J.C. Bose National Fellowship (SERB, Government of India) to AJ and, in part, by AJ’s personal funds.

## REFERENCES

Borash DJ and Ho GT 2001 Patterns of selection: stress resistance and energy storage in density-dependent populations of *Drosophila melanogaster*. J. Insect Physiol. 47 1349–1356

Case TJ 2000 An illustrated guide to theoretical ecology (Oxford: Oxford Univ. Press)

Dey S and Joshi A 2006 Stability via asynchrony in *Drosophila* metapopulations with low migration rates. Science 312 434–436

Dey S and Joshi A 2018 Two decades of *Drosophila* population dynamics: modelling, experiments, and implications; in Hand book of statistics, Vol. 39: Integrated population biology and modelling, Part A (eds) CR Rao and ASR Srinivasa Rao (Amsterdam and Oxford: Elsevier) pp. 275–312

Dey S, Bose J and Joshi A 2012 Adaptation to larval crowding in *Drosophila ananassae* leads to the evolution of population stability. Ecol. Evol. 2 941–951

Dey S, Prasad NG, Shakarad M and Joshi A 2008 Laboratory evolution of population stability in *Drosophila*: constancy and persistence do not necessarily coevolve. J. Anim. Ecol. 77 670–677

Ebenman B, Johansson A, Jonsson T and Wennergren U 1996 Evolution of stable population dynamics through natural selection. Proc. R. Soc. Lond. B 263 1145–1151

Ellner S and Turchin P 1995 Chaos in a noisy world: new methods and evidence from time-series analysis. Am. Nat. 145 343–375

Elton C 1927 Animal ecology (New York: Macmillan)

Gatto M 1993 The evolutionary optimality of oscillatory and chaotic dynamics in simple population models. Theor. Pop. Biol. 43 310–336

Gilpin ME, Case TJ and Ayala FJ 1976 θ-selection. Math. Biosci. 32 131–139

Grimm V and Wissel C 1997 Babel, or the ecological stability discussions: an inventory and analysis of terminology and a guide for avoiding confusion. Oecologia 109 323–334

Hansen TF 1992 Evolution of stability parameters in single-species population models: stability or chaos? Theor. Pop. Biol. 42 199–217

Hassell MP, Lawton JH and May RM 1976 Pattern of dynamical behaviour in single species populations. J. Anim. Ecol. 45 471–486

Joshi A 2022 Nine things to keep in mind about mathematical modelling in ecology and evolution. J. Biosci. 47 19

Joshi A and Mueller LD 1988 Evolution of higher feeding rate in *Drosophila* due to density-dependent natural selection. Evolution 42 1090–1092

Joshi A and Mueller LD 1996 Density-dependent natural selection in *Drosophila*, trade-offs between larval food acquisition and utilization. Evol. Ecol. 10 463–474

Joshi A, Prasad NG and Shakarad M 2001 *K*-selection, *α*-selection, effectiveness, and tolerance in competition: density-dependent selection revisited. J. Genet. 80 63–75

Kingsland SE 1995 Modeling nature, 2nd Edn (Chicago: University of Chicago Press)

Luckinbill LS 1978 r- and K-selection in experimental populations of Escherichia coli.

Luckinbill LS 1979 Selection and the *r–K* continuum in experimental populations of protozoa. Am. Nat. 113 427–437

MacArthur RH 1962 Some generalized theorems of natural selection. Proc. Natl. Acad. Sci. USA 48 1893–1897

MacArthur RH and Wilson EO 1967 The theory of island biogeography (New Jersey: Princeton University Press)

May RM 1974 Biological populations with non-overlapping generations: stable points, stable cycles and chaos. Science 186 645–647

Mueller L, Graves J and Rose MR 1993 Interactions between density-dependent and age-specific selection in *Drosophila melanogaster*. Func. Ecol. 7 469–479

Mueller LD 1988 Density-dependent population growth and natural selection in food limited environments: the *Drosophila* model. Am. Nat. 132 786–809

Mueller LD 1990 Density-dependent natural selection does not increase efficiency. Evol. Ecol. 4 290–297

Mueller LD and Ayala FJ 1981 a Dynamics of single species population growth: stability or chaos? Ecology 62:1148–1154.

Mueller LD and Huynh PT 1994 Ecological determinants of stability in model populations. Ecology 75 430–437

Mueller LD, Guo PZ and Ayala FJ 1991 Density dependent natural selection and trade-offs in life history traits. Science 253 433–435

Mueller LD, Nusbaum TJ and Rose MR 1995 The Gompertz model as a predictive tool in demography. Exp. Gerontol. 30 553–569

Mueller LD, Joshi A and Borash DJ 2000 Does population stability evolve? Ecology 81 1273– 1285

Muller LD 1997 Theoretical and empirical examination of density-dependent selection. The evolution of life-history traits. A critique of the theory and a review of the data. Annu. Rev. Ecol. Evol. Syst. 28 269–288

Nagarajan A, Natarajan SB, Jayaram M, Thammanna A, Chari S, Bose J, Jois SV and Joshi A 2016 Adaptation to larval crowding in *Drosophila ananassae* and *Drosophila nasuta nasuta*: increased larval competitive ability without increased larval feeding rate. J. Genet. 95 411–425

Prasad NG, Dey S, Shakarad M and Joshi A 2003 The evolution of population stability as a by-product of life-history evolution. Biol. Lett. 270 S84–S86

Reznick D, Bryant MJ and Bashey F 2002 *r*- and *K*-selection revisited: the role of population regulation in life history evolution. Ecology 83 1509–1520

Ricker WE 1954 Stock and recruitment. J. Fish. Res. Board Can. 11 559–623

Sah P, Salve JP and Dey S 2013 Stabilizing biological populations and metapopulations through Adaptive Limiter Control. J. Theor. Biol. 320 113–123

Santos M, Borash DJ, Joshi A, Bounlutay N and Mueller LD 1997 Density-dependent natural selection in *Drosophila*: evolution of growth rate and body size. Evolution 51 420–432

Sarangi M 2018 Ecological details mediate different paths to the evolution of larval competitive ability in *Drosophila*. PhD thesis, Jawaharlal Nehru Centre for Advanced Scientific Research, Bangalore

Sarangi M, Nagarajan A, Dey S, Bose J and Joshi A 2016 Evolution of increased larval competitive ability in *Drosophila melanogaster* without increased larval feeding rate. J. Genet. 95 491–503 Science 202 1201–1203

Sheeba V and Joshi A 1998 A test of simple models of population growth using data from very small populations of *Drosophila melanogaster*. Curr. Sci. 75 1406–1410

StatSoft 1995 Statistica Vol. I: general conventions and statistics 1. (Tulsa: StatSoft Inc.)

Stokes TK, Gurney WSC, Nisbet RM and Blythe SP 1988 Parameter evolution in a laboratory insect population. Theor. Popul. Biol. 34 248–265

Thomas WR, Pomerantz MJ and Gilpin ME 1980 Chaos, asymmetric growth and group selection for dynamical stability. Ecology 61 1312–1320

Tung S, Rajamani M, Joshi A and Dey S 2019 Complex interaction of resource availability, life-history and demography determines the dynamics and stability of stage-structured populations. J. Theor. Biol. 460 1–12

Turelli M and Petry D 1980 Density-dependent selection in a random environment: an evolutionary process that can maintain stable population dynamics. Proc. Natl. Acad. Sci. U.S.A. 77 7501–7505

Vaidya GP 2013 Dynamics of crowded populations of *Drosophila melanogaster*. MS thesis, Jawaharlal Nehru Centre for Advanced Scientific Research, Bangalore

